# MORC2, a sentinel of HIV-1 expression through HUSH-Mediated Epigenetic Silencing and RNA Degradation

**DOI:** 10.1101/2023.03.29.534756

**Authors:** Angélique Lasserre, Sébastien Marie, Marina Morel, Michael M. Martin, Alexandre Legrand, Virginie Vauthier, Andrea Cimarelli, Lucie Etienne, Florence Margottin-Goguet, Roy Matkovic

**Author notes:** co-first authors (Angélique Lasserre, Sébastien Marie).

## Abstract

The HUSH complex (composed of TASOR, MPP8 and periphilin) represses HIV-1 expression by inducing both propagation of repressive epigenetic marks and degradation of the nascent transcript. Vpx from HIV-2, and Vpr proteins from some simian lentiviruses (SIVs), antagonize HUSH, thereby increasing proviral expression. The chromatin-remodeler MORC2 protein plays a critical role in the epigenetic silencing of host genes by HUSH. Here, we deciphered the role of MORC2 in HIV-1 silencing. We show that MORC2, in contrast to HUSH components, presents strong signatures of positive selection during primate evolution. However, while HUSH is degraded upon HIV-2 infection in a Vpx-dependent manner, MORC2 levels are rather increased, due to the loss of the HUSH-mediated repression. Our results suggest that lentiviral proteins from the Vpr/Vpx family have not driven primate MORC2 evolution. Our findings indicate that MORC2 negatively regulates HIV-1 provirus expression. Mechanistically, we show that MORC2 is recruited to the integrated HIV-1 provirus locus and is required for TASOR-mediated post-transcriptional HIV-1 silencing suggesting that MORC2 sets the stage for the HUSH-mediated HIV-1 nascent RNA degradation. We demonstrate that reducing MORC2 levels diminishes provirus silencing in both monoclonal and polyclonal cellular models of HIV-1 latency. These results suggest that MORC2 has evolved adaptations during primate diversification, possibly in response to challenges posed by DNA pathogens or retroelement integration into the host genome.

## Introduction

Human Immunodeficiency viruses type 1 and 2 (HIV-1 and -2) are responsible for Acquired Immunodeficiency Syndrome (AIDS). Their eradication faces a major obstacle that is viral persistence in reservoir cells, despite antiretroviral treatment. HIV latency refers to silent viruses, which can produce infectious particles in case the treatment is interrupted. While HIV has long been thought to be transcriptionally inactive in latently infected cells, recent works now show that transcription can occur in these cells and that latency may result in co-/post-transcriptional repression of HIV-1 expression (1).

TASOR, MPP8 and Periphilin-1 have recently been shown to be major regulators of HIV-1 expression by forming the HUman Silencing Hub (HUSH) complex component. The mechanism by which HUSH recognizes and silences HIV is still poorly understood. MPP8 is suggested to tether the HUSH complex to the targeted chromatin locus through binding to H3K9me3 marks *via* its chromodomain, so that HUSH induces the propagation of H3K9me3 with the help of the histone methyltransferase SETDB1 (2). In the aim to understand how HUSH might regulate mechanistically HIV-1 silencing, we discovered that the HUSH core member TASOR binds both the elongating RNA Polymerase 2 and the nascent transcript and recruit RNA degradation/destabilisation factors such has the RNA exosome and CNOT1 leading to HIV-1 silencing at both the transcriptional and post-transcriptional steps(3). Cooperation between HUSH and the same RNA decay machineries was then confirmed to be also operating on host transposable elements in mouse ESC lines (4).

On one hand, HUSH represses HIV and lentiviral expression, while on the other hand, HUSH is counteracted by lentiviral proteins, including Vpx from HIV-2 and Vpr from several non-human primate lentiviruses (5–7). From an evolutionary standpoint, viral antagonism of the host antiviral factors, often triggers a strong pressure on these host proteins, and only those able to escape the viral antagonism will be selected along evolution. In turn, the viral protein is also submitted to a selective pressure to maintain antagonism. Over millions of years of coevolution, conflicts between hosts and pathogens have left marks on the protein-coding sequences of orthologous genes (genes in different host species that originated from a common ancestor). These conflicts are then evident in the selection of advantageous mutations in the host, which is indicated by a higher rate of non-synonymous substitutions (dN) compared to the rate of synonymous substitutions (dS), particularly at the direct virus-host protein molecular interface (8–10). Such signatures of positive selection are notably present in the HIV restriction factors SAMHD1, APOBEC3G or Tetherin/BST-2 – all three antagonized by specific lentiviral proteins (reviewed in (9)). Despite HUSH being degraded by lentiviral lineage-specific proteins, HUSH subunits are not characterized by strong positively selected sites (2). However, signs of the molecular conflict between HUSH and lentiviral proteins may also be spotted through protein diversification, like generation of multiple isoforms in HUSH members (ie: Periphilin-1) (5) or through identification of positive selection in another cellular partner of HUSH that may serve as a molecular bridge between HUSH and lineage-specific Vpr/Vpx.

Following the discovery of HUSH, MORC2 (Microrchidia family CW-type zing finger 2) was identified as a protein critical for HUSH-mediated epigenetic silencing of host genes (11). Alterations in the MORC2 gene are associated with the Charcot Marie Tooth disease (12–16). MORC2 belongs to the MORC ATPase superfamily (MORC1-4 and the divergent SMCHD1), which has been implicated in epigenetic silencing in diverse organisms (17–19). It is a chromatin-remodelling protein, which contains, alike other proteins from the family, an N-terminal gyrase, Hsp90, histidine kinase and MutL (GHKL)-type ATPase domain and a CW-type zinc finger histone recognition module (17). In addition, MORC2 possesses a C-terminal chromo-like domain (CD domain), which is a domain involved in the recognition of methylated peptides in histones and non-histone proteins (17). MORC2 is also involved in the DNA damage response contributing to a PARP-1-dependent repair pathway (20) and in the cell cycle control through an acetylation-dependent mechanism (21). The HUSH complex has been proposed to recruit MORC2 to specific loci, where the MORC2 ATPase and CW domains are required for HUSH silencing (11). In contrast, the CD domain does not appear important for MORC2 repression activity (11). Because chromatin remodelling *in vitro* relies on the ATPase activity (22), it was suggested that MORC2 remodelling activity is required to alter chromatin architecture and to promote gene silencing by HUSH (11). Indeed, loss of MORC2 resulted in the loss of H3K9me3 marks, chromatin decompaction and loss of transcriptional silencing at HUSH target sites (11). Apart its broad distribution across H3K9me3-marked heterochromatin, MORC2 was also detected at CpG-rich transcription start sites (TSSs) independently from HUSH (11).

Whether MORC2 is involved in the silencing of the HIV-1 provirus has not been investigated yet. We show here that MORC2 presents the features of a restriction factor with strong signatures of positive selection as well as antiviral activity. We also provide lines of evidence showing that MORC2 is required for HUSH-mediated HIV-1 silencing and participate in the transcriptional and post-transcriptional silencing of HIV-1 expression.

## Results

The presence of strong signatures of positive selection during primate evolution is a frequent characteristic of lentiviral restriction factors (9). When MORC2 was identified as an essential protein for HUSH repression, we wondered whether MORC2 has been evolving under positive selection in primates, in contrast to HUSH components (5). To address this possibility, we analysed MORC2 orthologous sequences from 25 species of primates, reconstructing its evolutionary history over 60-80 million years of divergence. The topology of the phylogenetic tree reconstructed from the primate MORC2 coding sequences was concordant with the primate species tree (Fig. 1A). To determine if MORC2 has evolved under positive selection, we performed several positive selection analyses available in the Detection of Genetic INNovation (DGINN) pipeline and from the HYPHY package (see Methods)(23). We first found strong evidence of positive selection at the gene-wide level in primate MORC2 (Fig. 1B). Furthermore, site-specific analyses identified at least nine sites undergoing rapid evolution, notably in the C-terminal part of the protein (Fig. 1C-D). The presence of two clusters of positively selected residues within the disordered regions surrounding the chromo-like domain (CD) may indicate viral specificity domains (24) (Fig. 1E-F). These results suggest that MORC2 might have been involved in a molecular conflict along evolution and that it might play a unique role in HUSH-mediated HIV restriction.

**Figure 1:**
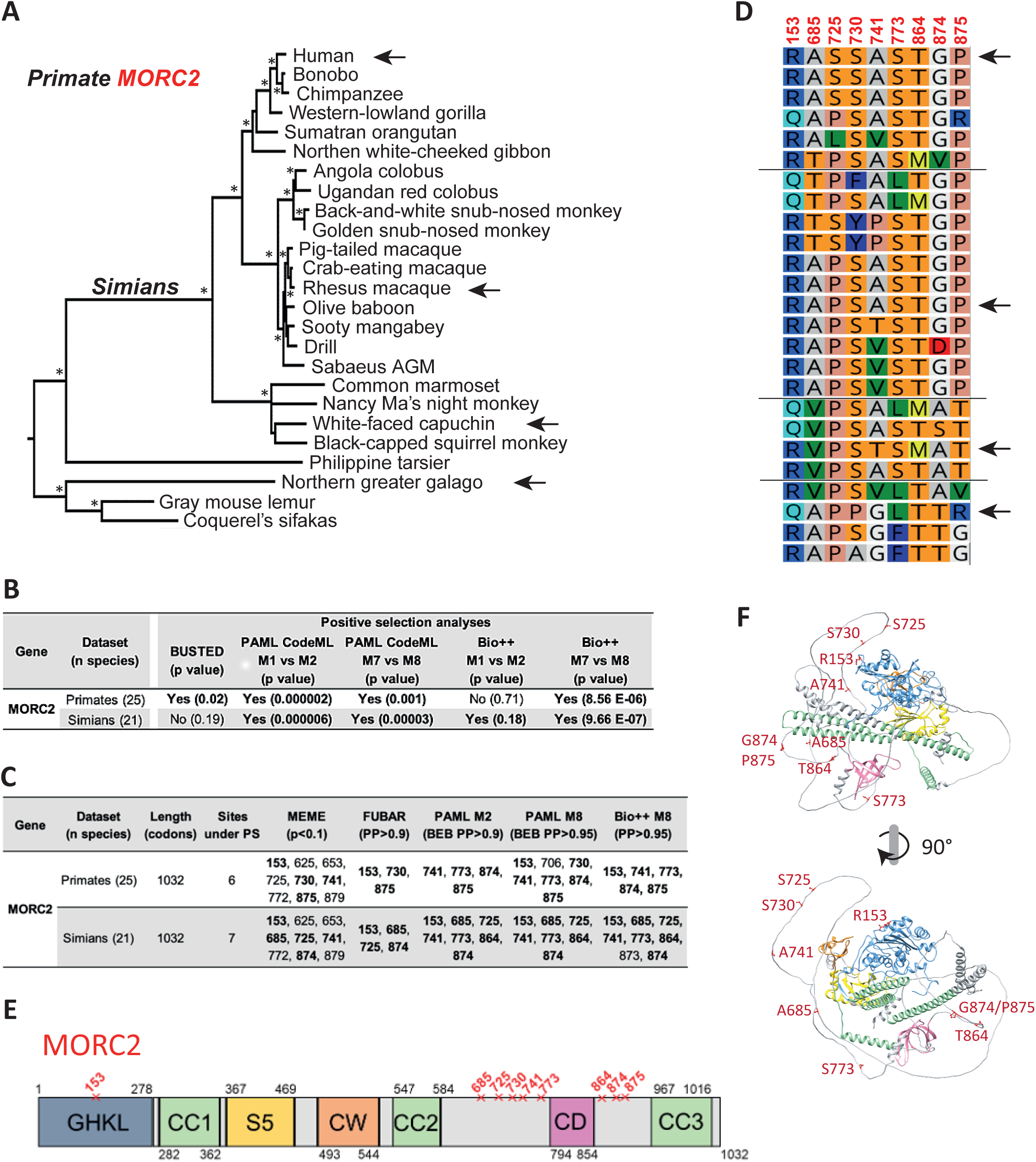
MORC2 presents signatures of positive selection during primate evolution and anti-lentiviral/HIV activity. **A. Phylogenetic analysis of primate MORC2.** The tree was built with PhyML and statistical support is from 1,000 bootstrap replicates (asterisks correspond to bootstrap values >0.75). The scale bar indicates the number of substitutions per site. Four black arrows correspond to the orthologous MORC2s that were subsequently tested in functional assays. **B. Evidence of positive selection during primate MORC2 evolution.** Analyses were performed with the DGINN pipeline, using the primate and the simian codon alignments at the gene-wide level (from 25 and 21 species, respectively). We used five methods to assess for positive selection: BUSTED from HYPHY, as well as two analyses from PAML CodeML and from Bio++ (see methods). P values that support a model allowing for positive selection are shown in parentheses. **C. Sites under positive selection in primate MORC2.** Site-specific positive selection analyses were performed using the DGINN pipeline and the Datamonkey server: MEME and FUBAR, BEB from PAML Codeml M2 and M8, and PP from Bio++ M8. Only the sites above the indicated “statistically significant cut-off” are shown. In bold are the sites identified by several methods. **D. Amino acid alignment of the sites under positive selection.** The species correspond to the panel A. Coloring is according to RasMol (Geneious, Biomatters). **E. Schematic of MORC2 with its functional domains and the herein identified sites under positive selection.** The ATPase module contains the ATPase domain, the coiled-coil domain 1 and the S5 domain. CC: Coiled-coil domain, CW: CW domain with zinc finger, CD: Chromodomain. Sites under positive selection from primate and simian analyses are indicated by red crosses. **F. Predictive 3D structure representation of MORC2 domains and sites under positive selections.** AlphaFold structure prediction of human MORC2. Functional domains are coloured following the panel E colour code. Atoms of the residues under positive selection are coloured in red and represented with sticks. Corresponding coordinates are labelled in red.

We therefore investigated whether MORC2 restricts an HIV-1-derived virus. HeLa cells were first infected with a VSVg-pseudo-typed (VSVg for Vesicular Stomatis Virus G envelope protein) non-replicative HIV-1 retrovirus that packaged a CMV-Luc (Luciferase) cassette flanked by the two HIV-1 LTRs. After reverse transcription, this virus integrates the 5’-LTR-CMV-Luc-3’-LTR construct into the host genome and luciferase transcription/expression is driven by the strong CMV promoter. This construct was previously identified as a good readout of HUSH activity (5) as we have shown that HUSH activity was linked to active transcription elongation (3). We found that siRNA-mediated MORC2 depletion increased Luc expression, while the overexpression of WT MORC2 triggered the opposite (Sup Fig. 1A, and 1B). The MORC2 point mutant (D68A), which impairs ATPase activity and chromatin remodelling (11, 22), abolished its repressive activity, suggesting that MORC2-mediated CMV-Luc repression relies on its ability to hydrolyze ATP and perhaps to its chromatin remodelling activities (Sup Fig. 1B). Repression was also obtained with a SIVmac-derived virus, suggesting MORC2 has a broad anti-lentiviral activity (Sup Fig. 1A, right). To point out a HIV-1 LTR-dependent effect, we further used a Hela cellular system in which the luciferase gene reporter is directly under the control of the HIV-1 LTR promoter integrated in the host genome and deleted of the TAR element sequence that blocks RNA polymerase II (RNAPII) elongation. Thus, only expression from the integrated construct is analysed and productive transcription can be studied independently of RNA splicing and RNAPII pausing (3). MORC2 depletion led to an increase in HIV-1 LTR-driven expression, while the expression of WT MORC2, but not MORC2 D68A, decreased luciferase expression (Fig. 2A and 2B). Notably, the R252W substitution, the most common alteration affecting MORC2 in Charcot-Marie-Tooth patients, which was proposed to hyper-activate HUSH-silencing of host genes in neuronal cells was also effective in HIV-1 expression repression (11). Furthermore, MORC2 deleted of its CD chromodomain still had a repressive effect, while MORC2 lost its activity when the CC2 domain was removed, as expected from a study using a different promoter (11).

**Figure 2:**
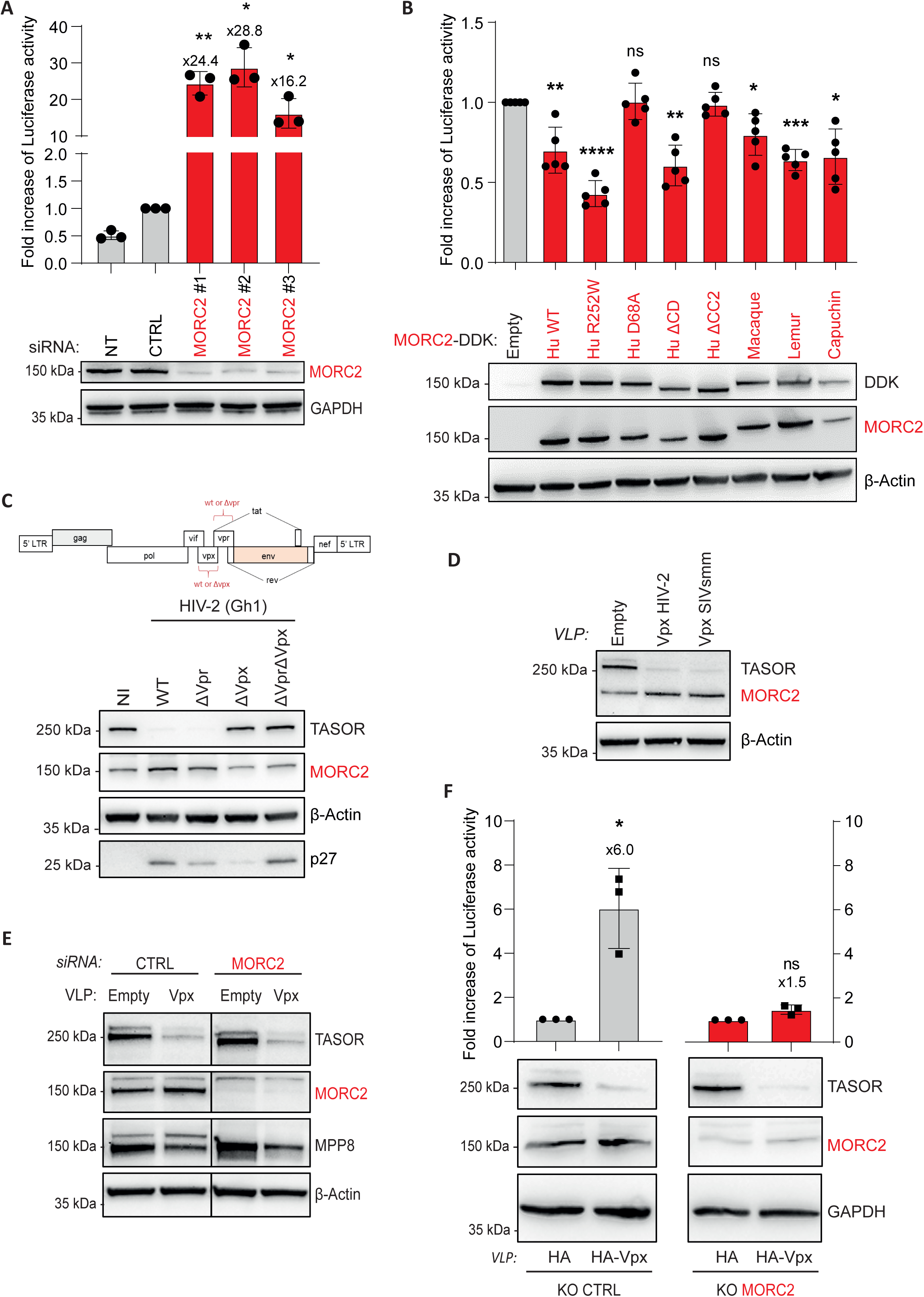
MORC2 is an antiviral protein upregulated by HIV-2 Vpx. **A. MORC2 inhibition upregulates HIV-1 5’ LTR expression.** HeLa HIV-1 LTR ΔTAR-Luc cells were transfected with 3 different siRNAs that antagonize specifically MORC2 mRNAs. The Luciferase activity was measured, normalized to BCA for each condition and to the siRNA control condition 3 days post transfection. (*n* = 3; each replicate is presented along with the mean values and the SD. Lognormality has been verified with a Shapiro-Wilk test prior to a One sample t test with a theoretical mean of 1. \**p* < 0.05 \*\**p* < 0.01) **B. Overexpression of Human and Non-Human primate MORC2 orthologs silence HIV-1 LTR expression in Human cells.** HeLa HIV-1 LTR ΔTAR-Luc cells were transfected with cDNAs encoding Human (Hu), or Macaque, Lemur, Capuchin MORC2-DDK. The Luciferase activity was measured, normalized to BCA for each condition and to the Empty pCMV6-DDK condition 2 days post-transfection. (*n* = 5; each replicate is presented along with the mean values and the SD. Lognormality has been verified with a Shapiro-Wilk test prior to a One sample t test with a theoretical mean of 1. \**p* < 0.05 \*\**p* < 0.01 \*\*\**p* < 0.001 \*\*\*\**p* < 0.0001) **C. TASOR degradation upon HIV-2 infection is accompanied by a slight increase in MORC2 levels**. Jurkat T cells were non-infected (NI) or infected with complete HIV-2 viruses (WT), or HIV-2 viruses deleted of either Vpr (ΔR), Vpx (ΔX) or both (ΔRΔX). Levels of the indicated proteins were analyzed by western blot. **D. MORC2 levels are upregulated in the presence of Vpx from HIV-2 or from SIVsmm.** HeLa cells were treated by HIV-2 or SIVsmm Vpx-containing VLPs. Cells were lysed 24h post-transduction and Western-Blot was performed. **E. MORC2 is not required for Vpx-mediated HUSM member TASOR degradation.** HeLa cells were treated by siRNA against MORC2, then transduced by VLP containing Vpx. **F. MORC2 is required for TASOR-mediated LTR-driven Luciferase silencing.** VLPs containing HIV-2 HA-Vpx or HA (empty) were transduced for 24h into HeLa HIV-1 LTR ΔTAR-Luc cells knocked out (KO) for MORC2 or Mock (Ctrl). Luciferase activity was measured 24h after VLP transduction, and normalized to BCA. Data is represented as fold increase to the empty VLP condition. (*n* = 3; each replicate is presented along with the mean values and the SD. Lognormality has been verified with a Shapiro-Wilk test prior to a One sample t test with a theoretical mean of 1. \**p* < 0.05)

We further addressed the possibility that MORC2 from primate orthologs could have different abilities to repress HIV-1, as species-specificity may emerge from rapidly evolving restriction factors. By expressing four orthologous MORC2 proteins from highly divergent primate species, whose differences have led to the designation of positive selection sites, we found that they similarly repressed viral expression in our two experimental systems, infection by a HIV-1-LTR-CMV virus (Sup Fig. 1B) or expression driven by the integrated LTR promoter (Fig. 2B). These results show that MORC2 retains a well-conserved repressive activity. Furthermore, our heterologous virus-host assay, which included MORC2 from four primate species with diverse amino acids at the positively selected sites (Fig. 1D), suggests that positive selection of MORC2 is not linked to the restrictive effector function against the HIV-1-derived virus. Together, our results suggest that primate MORC2 ortholog proteins retain a well conserved repressive activity from the LTR-mediated integrated CMV promoter or directly from the LTR promoter in Human cells.

Given that HUSH recruits MORC2 and is inactivated by divergent lentiviral Vpr or Vpx proteins and that both HIV-2/SIVsmm Vpx and HIV-1 Vpr have been reported to enhance LTR-driven transcription(5, 25–28), we investigated whether HIV-2/SIVsmm Vpx or HIV-1 Vpr could antagonize MORC2 or use it for their functions. Unlike HUSH, MORC2 was not degraded following infection by a complete WT HIV-2 virus; instead, it was slightly stabilized, in a Vpx-dependent manner (Fig. 2C). Accordingly, Vpx-containing Viral-Like pseudo-Particles (VLPs) induced TASOR degradation and MORC2 stabilization as reported in (7) (Fig. 2D). MORC2 mRNA levels were increased with TASOR depletion, suggesting that MORC2 stabilization results from a transcriptional effect, in line with previous reports (3, 7, 11) (Sup. Fig. 2A). Regarding HIV-1, MORC2 levels were unaffected following infection, while levels of the HLTF DNA translocase were slightly reduced in a Vpr-dependent manner as expected (30) (Sup. Fig. 2B). Further testing of a set of divergent Vpr and Vpx lentiviral proteins revealed that none of them could induce MORC2 destabilization (Sup. Fig. 2C).

Considering that positive selection sites may indicate a surface of interaction with viral proteins (8–10), we also investigated whether MORC2 could link HUSH and Vpx to induce HUSH degradation. MORC2 depletion by siRNA did not affect Vpx-mediated HUSH degradation (Fig. 2E, compare 3,4 to 1,2), or Vpr-mediated HLTF degradation (Sup. Fig. 2D, compare lanes 4,5 to 1,2) (29, 30). Similarly, Vpx-mediated TASOR degradation was not impaired in MORC2 knockout cells (Fig. 2F). Nonetheless, in these cells, the ability of Vpx to increase LTR-driven expression was significantly inhibited, indicating that TASOR is repressing HIV-1 expression with the help of MORC2 as it is the case for host genes (11). Altogether, these results suggest that HIV-2 Vpx and HIV-1 Vpr do not induce MORC2 degradation, nor do they use MORC2 to induce the degradation of HUSH or HLTF, respectively. However, this does not exclude the possibility that Vpx or Vpr could inactivate MORC2 in any way other than degradation. In addition, MORC2 is essential for HUSH-mediated silencing of HIV-1 LTR-driven expression.

Next, we explored the role of MORC2 in more relevant cell models of HIV infection. We used the J-Lat A1 monoclonal model of HIV-1 latency, derived from the Jurkat T cell line, which contains an HIV-1 LTR-*tat*-*IRES*-*gfp-*LTR minigenome stably integrated at a unique site and epigenetically silenced (31). An increase in the percentage of GFP-positive cells is indicative of viral reactivation from the latent state. The depletion of MORC2 by CRISPR/Cas9 led to an increase of GFP expression in these cells (Fig. 3A). Cells were FACS-sorted according to GFP expression (no, low or high GFP expression) and put back in culture for two weeks. Western blot analyses confirmed that the more MORC2 was depleted, the more GFP was expressed (Fig. 3A right, Western Blot). Secondly, we used another Jurkat model of HIV-1 latency we previously generated (3), in which the latent proviruses retain complete LTRs, Tat and Rev, but has a frameshift mutation in Env, nef growth factor receptor (ngfr) in place of Nef and GFP in place of Gag, Pol, Vif and Vpr (7). In this polyclonal latency model, latently infected cells were not selected for a particular integration site but represent a population with diverse numbers of integration sites and at different genomic loci. Depletion of MORC2 led to an increase of the eGFP expression, with or without TNFα−mediated HIV-1 LTR transcription activation (Fig. 3B). Altogether, MORC2 appears as sentinel to keep HIV-1 silent.

**Figure 3:**
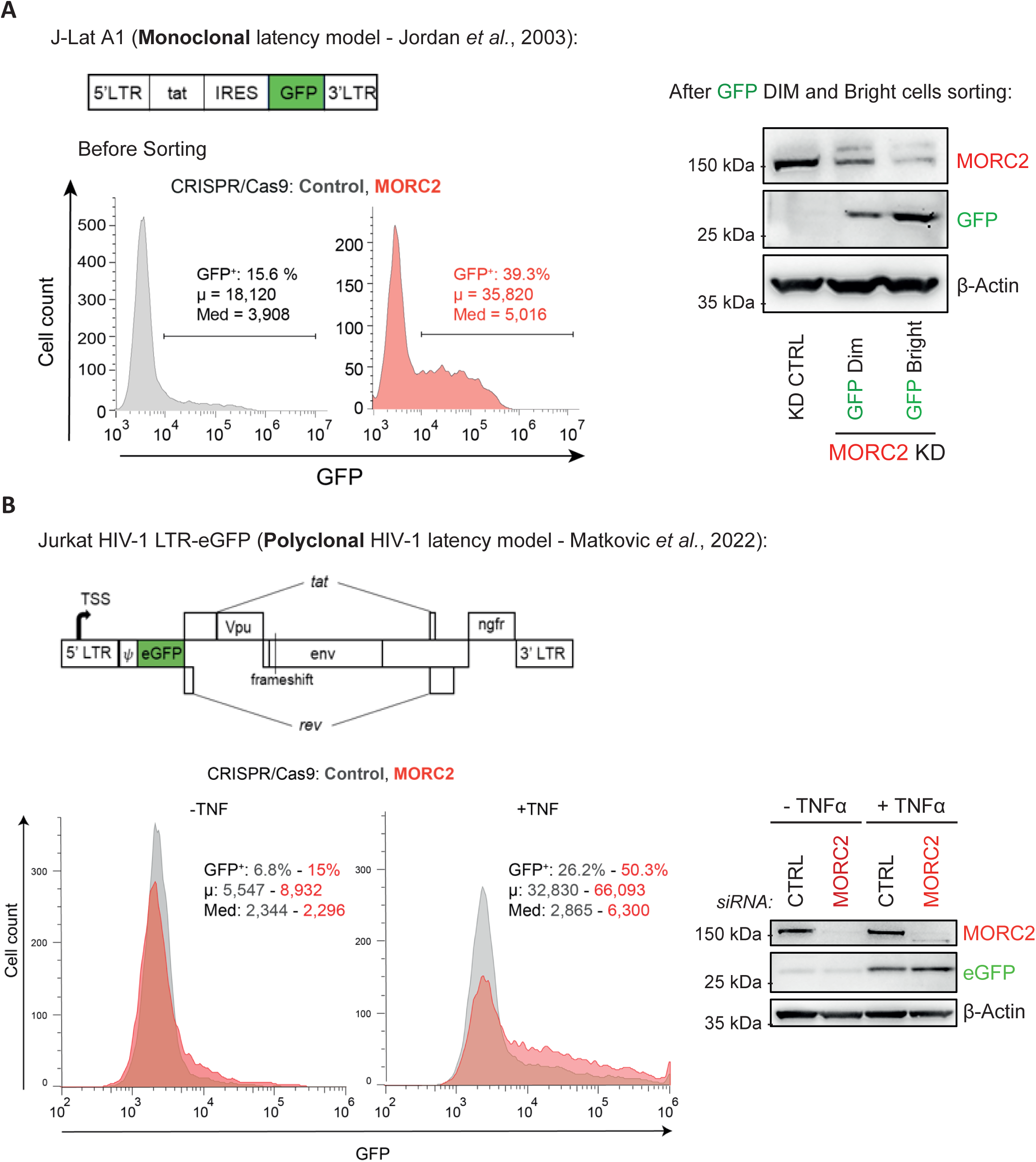
MORC2 presents a repressive effect in two models of HIV-1 latency. **A. MORC2 represses HIV-1 expression in the J-Lat A1 model of latency.** *Left*. J-Lat A1 cells, harboring one copy of integrated latent LTR-tat-*IRES*-GFP-LTR construct, were depleted of MORC2 by CRISPR/Cas9, which leads to an increase of the percentage of GFP-positive cells (reactivation of the virus analyzed by cytometry). *Right.* MORC2 depletion accompanies viral reactivation gradually. J-Lat A1 Cells were FACS-sorted according to GFP expression. More LTR-driven GFP expression was high, less MORC2 was expressed. **B. MORC2 represses HIV-1 LTR-driven eGFP expression in a polyclonal CD4+ model of HIV-1 latency.** Jurkat T cells latently infected by HIV-1-eGFP (the viral genome is schematically shown above) were depleted of MORC2 by CRISPR/Cas9, which leads to enhanced HIV-1 expression without and with TNFα.

Next, we wanted to better decipher the mechanisms by which MORC2 does negatively control HIV-1 expression as HUSH needs MORC2 to silence gene and HIV-1 expression. We previously discovered that TASOR was acting at the co-/post-transcriptional level to induce the degradation of the HIV-1 LTR-driven nascent RNA by recruiting and cooperating with the nuclear CNOT1 and the RNA exosome containing complexes (3). Thus, we performed a *Nuclear Run On* experiment in the HIV-1-LTRΔTAR-Luc system, in which TASOR has a negative impact on the LTR-driven RNA stability, to distinguish MORC2 effects i) on nascent luciferase transcript levels, labelled with BrUTP, and ii) on total luciferase mRNA levels. As expected, TASOR down-regulation poorly impacted the levels of nascent transcripts in this system, while it triggered a 2.3-fold increase of steady-state Luc RNA levels, in agreement with a role of TASOR in the turnover of LTR-driven transcripts in this system (Fig. 4A)(3). MORC2 depletion induced a slight increase in HIV-1 LTR nascent transcript levels (1.8 fold) and a better accumulation of HIV-1 LTR-driven RNA at the total level (3.7 fold), suggesting that MORC2 depletion induces HIV-1 LTR-driven RNA stabilization (Fig. 4A). Because MORC2 seems to induce the HIV-1 LTR-driven RNA destabilization, we investigated whether MORC2 was able to cooperate with CNOT1, as this TASOR partner does the same on the nascent LTR-driven Luc transcript in this HIV-1-LTRΔTAR-Luc system (3). The siRNA-mediated co-silencing of MORC2 and CNOT1 expressions induced a strong synergistic effect on the LTR-driven Luciferase expression, like the co-depletion of TASOR and CNOT1 (Fig. 4B)(3). This suggests that MORC2 and CNOT1 use different pathways that are connected to better repress HIV-1 expression. The triple depletion of TASOR, MORC2 and CNOT1 resulted in a greater synergistic effect, highlighting a potential cooperation between the three proteins to repress the HIV-1 LTR (Fig. 4B). In agreement with a cooperation, we found that endogenous MORC2 interacts with CNOT1 in nuclei of HeLa cells (Fig. 4C). This interaction was reduced by RNAse A treatment, suggesting that RNAs play a role in the association between MORC2 and CNOT1 (Fig. 4C). Furthermore, following immunoprecipitations of the nuclear CNOT1 in HeLa cells that were CRISPR/Cas9-KO for TASOR or MORC2 expression, we show that MORC2 was able to interact with CNOT1 in absence of TASOR, and TASOR with CNOT1 in absence of MORC2 (Sup. Fig. 3A). These results suggest that TASOR and MORC2 may associate with CNOT1 in an independent-manner forming different repressive CNOT1-containing complexes. We previously showed that TASOR was also interacting and cooperating with the RNA exosome containing complexes like MTR4 and the PAXT complex subunit ZFC3H1 (3). We then wondered whether MORC2 was able to interact with these same TASOR partners involved in nuclear RNA degradation. Through exogenously expressed MORC2 immunoprecipitation, we retrieved CNOT1, but also MTR4 and ZFC3H1 (Fig. 4D). These proteins were also found in association with endogenous MORC2 (Sup. Fig. 3B). Despite CNOT1 interacts with CD domain-containing proteins such as MPP8 (32), the removal of MORC2 CD domain did not impact its interaction with CNOT1, MTR4 or ZFC3H1 (Fig. 4D). It is worthy to note that while MORC2 depletion does reduce the silencing effect of TASOR in the HeLa HIV-1-LTRΔTAR-Luc system, suggesting that MORC2 sets the stage for the co-/post-transcriptional control of HIV-1 expression mediated by TASOR, its absence does stimulate the CNOT1 silencing pathway (Fig. 4E). This shows that TASOR/MORC2, on one hand, and CNOT1 on the other, use different pathways, but still connected, to repress HIV-1 expression. Altogether, MORC2 appears to repress HIV-1 LTR expression at the transcriptional level, and is found to be regulating directly or indirectly the stability of the LTR-driven transcript by interacting notably with TASOR and RNA degradation proteins. To understand whether the repressive functions of MORC2 on HIV-1 LTR expression are direct, we performed Chromatin-immunoprecipitation experiments coupled to qPCR in the HIV-1-LTR-ΔTAR-Luc system. While we do detect MORC2 on the TAF7, ZNF224, TUG1 genes (Fig. 5) as expected from ChIP-seq data in K562 cells (60) (Sup. Fig. 5) we show that MORC2 is also associated with both the HIV 5’LTR promoter and the Luc coding sequence, in agreement with its role in assisting TASOR-mediated Gene/HIV-1 repression during transcription (3, 33)(Fig. 5).

**Figure 4:**
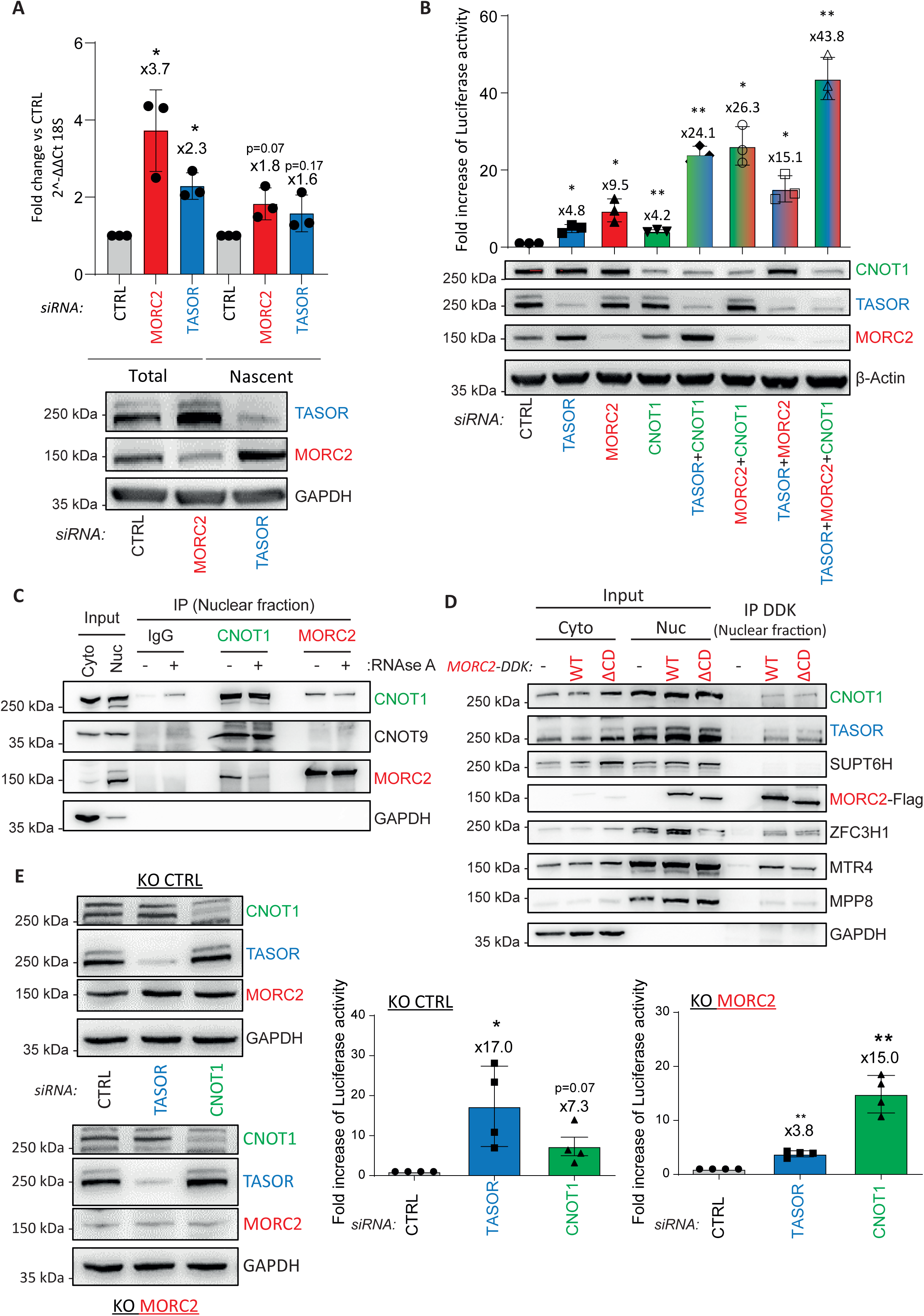
MORC2 induces HIV-1 LTR-driven transcripts destabilization and cooperates with CNOT1. **A. MORC2 negatively impacts LTR-driven transcripts at both transcriptional and post-transcriptional levels**. HeLa HIV-1 LTR ΔTAR-Luc cells were transfected for 72 h with Ctrl or MORC2 siRNA. Nuclear Run On experiments were performed to analyze the effect of MORC2 depletion on the level of nascent and total RNAs from the LTR *(Top).* Western Blot was performed to check the depletion of MORC2 and TASOR *(Bottom)*. (*n* = 3; each replicate is presented along with the mean values and the SD. Lognormality has been verified with a Shapiro-Wilk test prior to a One sample t test with a theoretical mean of 1. \**p* < 0.05) **B. MORC2, TASOR and CNOT1 cooperate to repress expression from the HIV-1 LTR.** HeLa HIV-1 LTR ΔTAR-Luc cells were transfected for 72 h with one or two or three siRNAs as indicated. Luc activity *(Left)* was measured (*n* = 3; each replicate is presented along with the mean values and the SD. Lognormality has been verified with a Shapiro-Wilk test prior to a One sample t test with a theoretical mean of 1. \**p* < 0.05 \*\**p* < 0.01) and protein depletion was checked by Wester-Blot. C. Transcripts facilitate the interaction between MORC2 and CNOT1. Nuclear fraction of HeLa cells was isolated and incubated with RNAse A or not for 30 minutes at room temperature. Then, endogenous CNOT1 or MORC2 were immunoprecipitated with specific antibodies as indicated and Western-Blot performed. The cytoplasmic GAPDH protein is a control of the nucleus-cytoplasm fractionation. **D. The MORC2 Chromodomain is not involved in the interaction with CNOT1, and PAXT associated proteins MTR4/ZFC3H1.** MORC2-DDK WT and MORC2-DDK ΔCD encoding vectors and the corresponding empty vector were transfected in HeLa cells. After 48h, cells were lysed and the nuclear fraction was isolated. Anti-DDK immunoprecipitation on this nuclear fraction and Western Blot were performed. **E. MORC2 is required for TASOR- but not for CNOT1-mediated LTR-driven RNA destabilization.** HeLa HIV-1 LTR ΔTAR-Luc cells KO Ctrl or MORC2 were transfected with siRNA TASOR or CNOT1 for 72h. The Luciferase activity was measured, normalized to BCA for each condition and to the siRNA control condition. (*n* = 4; each replicate is presented along with the mean values and the SD. Lognormality has been verified with a Shapiro-Wilk test prior to a One sample t test with a theoretical mean of 1. \**p* < 0.05 \*\**p* < 0.01)

**Figure 5:**
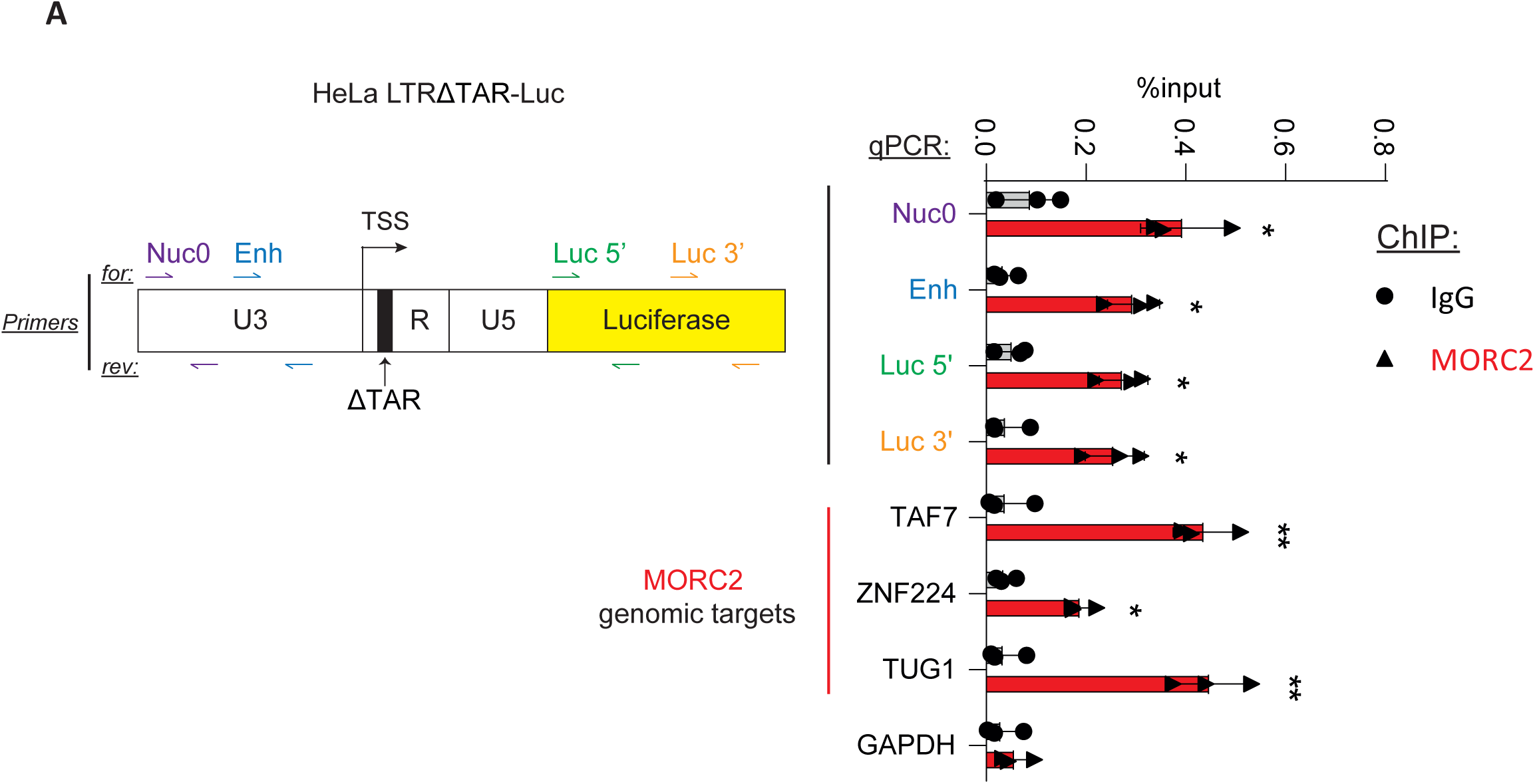
A. MORC2 binds the HIV-1 LTR promoter and its downstream coding sequence. Endogenous MORC2 occupancy was measured by ChIP-qPCR in HeLa HIV-1 LTR ΔTAR-Luc cells. TAF7, ZNF224, TUG1 genes are positive controls for MORC2 binding (see Sup. Fig 5). TSS stands for *Transcription Start Site.* (*n* = 3; each replicate is presented along with the mean values and the SD. Normality has been verified with an Anderson-Darling test and homoscedasticity with a Spearman test prior to a multiple two-tailed unpaired t test corrected with Šidák-Holm procedure. \**adj. p* < 0.05 \*\**adj. p* < 0.01)

## Discussion

MORC2 harbors signatures of positive selection along primate evolution, one of the well-defined characteristics of antiviral restriction factors. One possibility is to envision MORC2 as a component of the HUSH complex. Indeed, HUSH and MORC2 interact, and both proteins rely on each other for the silencing of incoming viruses, as shown here, or for the silencing of host genes (11). Therefore, HUSH associated with MORC2 could be seen as a single multi-subunit restriction factor that could provide different interfaces for viral attacks. In the case of the SAMHD1 restriction factor, Vpx/Vpr proteins from divergent viruses, despite their common origin and structural homology, target entirely different domains of the host protein in a species-specific manner (34–36). This difference in SAMHD1 recognition is evolutionarily dynamic and is further witnessed by sites of positive selection in both N-and C- terminal domains of the host protein (34, 37, 38). Analogously, HUSH-MORC2 could be targeted by different ways through their different subunits: HUSH by Vpx and MORC2 potentially by another lentiviral protein. The positively selected residues in MORC2 form two clusters in the disordered regions, which are also known to be frequent targets of positive selection (39, 40). These regions could constitute interfaces for binding and coupled folding with other proteins such as the HUSH complex, or viral proteins, where the subsequent acquired structure could vary depending on the partners. In any event, infection by HIV-1 or HIV-2 did not downregulate MORC2 expression, suggesting that MORC2 is not degraded by any of the viral proteins encoded by these viruses. Since MORC2 is not degraded, we wondered whether MORC2 could be a direct adaptor recruited by Vpx or Vpr to induce the degradation of their targets, such as CBFβ is recruited by Vif to ensure APOBEC3G degradation(41, 42). Though, noteworthy, CBFβ is evolutionary and structurally well conserved with no obvious sites of positive selection in contrast to APOBEC3G (43–45). We found that MORC2 was dispensable for Vpx-mediated HUSH or Vpr-mediated HLTF degradation. Altogether, we could not find any evidence that MORC2 is targeted by Vpr or Vpx or another lentiviral protein and the question of whether MORC2 evolution has been driven by lentiviruses remains open.

HUSH and MORC2 could also be considered as distinct entities. Indeed, HUSH subunits and MORC2 differ in several ways: (i) MORC2 presents sites of positive selection, not the HUSH components, (ii) MORC2, in contrast to HUSH, is not targeted by Vpx to the degradation pathway, (iii) in addition, MORC2 was found at Transcription Start Sites (TSS) in the absence of HUSH on host genes (11). Regardless of whether MORC2 works with HUSH, MORC2’s sites of positive selection might result from the MORC2’s engagement in an evolutionary arms race with a wide variety of entities other than lentiviruses. Candidates are viruses replicating *via* a DNA intermediate, present in the nucleus and silenced by repressive epigenetic marks, such as those from the *Herpesviridae* family, like Herpes simplex virus-1 (HSV-1)(46) or Cytomegalovirus (CMV)(47), or from the *Hepadnaviridae* family, like Hepatitis B Virus (HBV) (48). Recently, the ICP0 protein encoded by HSV-1 has been identified as a viral antagonist of MORC3, another member of the MORC family also involved in gene silencing and regulation of chromatin structure (49–52), highlighting *Herpesviridae* antagonists as potential drivers of the evolution of MORCs. The molecular conflict might also involve young and potentially adapting retroelements that are under the control of HUSH (33, 53–56). Finally, MORC2 may also have evolved under the selection process underlying speciation and adaptation as is the case for several other transcription factors or chromatin-associated proteins (57). It is interesting to note that among the 9 amino acids that have been positively selected over evolution, 4 of them are serine/threonine residues that can be phosphorylated in particular conditions, and that might regulate either its function within particular complexes and its localization. Similarly, we previously showed that TASOR is phosphorylated during mitosis on its threonine 819 (58), which might impact its association with chromatin localized complexes. Phosphorylation of the MORC2 S725, S739, S773 and T864 residues might then modulate its function/localization at the chromatin level.

Ironically, while Vpx induces HUSH degradation, MORC2 is stabilized in the presence of Vpx, due to a transcriptional effect: MORC2 gene expression being repressed by HUSH, the degradation of HUSH by Vpx induces an increase of MORC2 mRNA levels and subsequently of MORC2 protein levels (23). This stabilization effect at the protein level, previously reported in (7), is also observed when cells are infected with a complete WT HIV-2 virus compared to the ΔVpx virus. Whether MORC2 could contribute to HIV-1 expression inhibition once HUSH is degraded remains an open question. In these circumstances, MORC2 might provide a feedback control mechanism by limiting access of the HIV-1 LTR to transcription factors once TASOR is depleted. Indeed, MORC2 recruitment to gene promoters and to TSS was described to be HUSH-independent (11). It would be further interesting to quantify the HIV-1 LTR methylation modification upon HUSH depletion, as MORC2 is involved in the repression of endogenous genes by inducing methylation of promoters through the recruitment of DNMT3A (59). In line with this idea, gene silencing by RNA-directed DNA methylation in *Arabidopsis thaliana* depends on MORC proteins (60). Our results in two HIV-1 models of latency further suggest that MORC2 could be involved in HIV latency. Strikingly, with a different mechanism, MORC3 was proposed as a gatekeeper of its own anti-viral function: by degrading MORC3, HSV-1 unleashes the second antiviral activity of MORC3, which is the de-repression of the *IFNB1* gene expression and, subsequently, induces interferon production (51). Whether MORC2 could act as an immune gatekeeper following the degradation of HUSH, in particular during HIV-2 infection, is possible.

How MORC2 operates mechanistically is still poorly understood. We used vectors expressing Luciferase under different promoters (CMV and HIV-1 LTRΔTAR) and demonstrated that the silencing mechanism of MORC2 is independent of the promoter type in these strongly transcribing systems. Additionally, TNFα-induced stimulation of HIV-1 transcription from the 5’LTR in our HIV-1 latent systems, highlighted MORC2’s repressive function, similar to TASOR (3). These results suggest that MORC2 may be working with HUSH to tune down the expression of transgenes, re-establish HIV-1 latency, during RNAPII elongation activity in a co- or post- transcriptional fashion. We also showed that MORC2, like TASOR, acts directly or indirectly at the post-transcriptional level, and is found in complex with CNOT1 in an RNA-dependent manner and to synergistically cooperate with the latter to repress LTR-driven transcription. MORC2 also interacts with proteins from the PAXT complex, ZFC3H1 and MTR4, further supporting the hypothesis of a role of MORC2 in RNA destabilization. These findings support the notion of MORC2’s role in RNA metabolism and its coordination with TASOR, highlighting an integrated process that connects epigenetic regulation and RNA metabolism to regulate gene expression. The precise silencing mechanisms employed by MORC2 and its interplay with the RNA degradation complexes is yet to be further specified. Whether MORC2 cooperates with CNOT1 once HUSH is depleted remains a possibility since MORC2 and CNOT1 still interact in the absence of TASOR. MORC2 might act at the post-transcriptional level independently of HUSH, as suggested for the close related MORC3 protein. These questions are of paramount importance since MORC2 appears as a candidate player in the persistence of the virus in HIV-infected individuals.

## Materials and Methods

### Sequence and evolutionary analyses

The sequences of primate orthologous genes of MORC2 were retrieved from publicly available databases using NCBI Blastn, UCSC Blat or Ensembl, with the human sequence as the query. For phylogenetic and positive selection analyses, we retrieved one coding sequence per primate species for a total of 25 species. For each gene, two datasets were independently analyzed: one included the entire primate evolutionary history (spanning approximately 87 million years of primate evolution) and one was restricted to the simian primate species (spanning approximately 40 million years; with 21 simian primate species included). Orthologous sequences were codon-aligned using Muscle v3.8(61) and the phylogenetic tree was built using PhyML v3(62) with a HKY+I+G model with 1,000-bootstrap replicates for statistical support.

To comprehensively determine the evolution of MORC2 in primates, we performed positive selection analyses. Maximum-likelihood tests to assay for positive selection were performed using three packages: HYPHY v2.3(63), PAML v4(64) and Bio++ v2.2.0, all available within the DGINN pipeline(23, 65). In HYPHY, we used the branch-site unrestricted statistical test for episodic diversification (BUSTED) method, which detects gene-wide evidence of episodic positive selection within a codon alignment. In PAML, we used the Codeml program with the corresponding gene tree inferred from PhyML as input. Parameters were checked using the M0 one ratio model and the sequence alignments were then fit to models that disallow (M1 and M7) or allow (M2 and M8) positive selection. Likelihood ratio tests (LRTs) were performed to compare model M1 versus M2 and M7 versus M8, using a χ2 test to derive P values. We finally used the Bio++ package to similarly test for evidence of positive selection (M1^NS^ versus M2 ^NS^ and M7 ^NS^ versus M8 ^NS^, as implemented in Bio++; NS for non-stationary models). To look more specifically for site-specific positive selection, we used the fast, unconstrained Bayesian approximation for inferring selection (FUBAR) method from HYPHY/Datamonkey. In Codeml and Bio++, we used Bayesian empirical Bayes (BEB) analyses and Bayesian posterior probabilities, respectively, from the M2 and M8 models to determine if any codon was under significant positive selection.

Protein structure representations were conducted using UCSF Chimera (The Regents) software. Protein structure predictions are from AlphaFold Protein Structure Database(66).

### Plasmids and transfection

MORC2 expression vectors pCMV6-myc-DDK have been purchased from Origene. pCMV6-MORC2-myc-DDK vector expresses MORC2 isoform 1 of 1032 amino acids (NCBI Reference Sequence: NP_001290185.1). MORC2-R252W, MORC2-D68A, MORC2-ΔCD and MORC2-ΔCC2 constructs were obtained by directed mutagenesis using Phusion High Fidelity DNA polymerase (Thermo-Fisher). The pLentiCRISPRv2-sgMORC2-Cas9 and the pLentiCRISPRv2-sgTASOR-Cas9 were obtained by subcloning the sgRNAs targeting respectively the 5^th^ and the first exon of MORC2 and TASOR, with the BsmBI enzyme. All primers used to generate these plasmids are listed in Table S1. The pCMV6-MORC2-myc-DDK plasmid encoding the MORC2 cDNA from *Macaca mulatta* (rhesus macaque, NCBI Reference Sequence: NP_001248528.1), *Otolemur garnettii* (Northern greater galago, NCBI Reference Sequence: XP_003803380.2) and *Cebus capucinus imitator* (white-faced capuchin, NCBI Reference Sequence: XP_017355058.1) have been synthesized by Genecust through gene editing strategy.

All plasmids are transfected in cells the CaCl2 method.

### Virus and VLP production

All Virus Like Particles (VLP) and most of the viruses were pseudo-typed with VSV-G protein and produced in 293 FT (Gift from Nicolas Manel) through the CaCl2 co-precipitation method. Cell culture medium was collected 48h after transfection and filtered through 0.45 μm pore filters. Viral particles or VLPs were concentrated by ultra-centrifugation on sucrose gradient.

Regarding VLP production with viral proteins expressed in trans, SIV3+ ΔVprΔVpx packaging vector (gift from N. Landau(67)) and a VSV-G vector were co-transfected with pAS1B-HA-Vpx HIV-2 Ghana or pAS1B-HA (empty) in order to produced VLP containing HA-tagged Vpx protein. In the same manner pPAX2 packaging HIV-1 were co-transfected with a VSV-G vector and pAS1B-HA-Vpr HIV-1 or pAS1B-HA (empty) in order to produce VLP containing HIV-1 Vpr protein. For VLP with viral proteins expressed in cis, we transfected the VSV-G vector, the SIV3+ ΔVprΔVpx packaging vector (as the empty condition) or SIV3+ WT (R+X+ condition)(68) in order to produce VLP containing all accessories proteins encoded by the SIV3+ vector.

To deliver the sgRNAs and Cas9 to knock out MORC2, TASOR or Mock (control): pPAX2 packaging vector was co-transfected along with pLentiCRISPRv2-sgTASOR-Cas9, or with pLentiCRISPRV2-sgMORC2-Cas9 or with pLentiCRISPRV2-Cas9 (control) for VLP production (with VSV-G).

To produce HIV-1 viral particles: pPAX2 packaging was co-transfected with the HIV-1 LTR-eGFP transfer vector, described in(7) and obtained from Addgene (plasmid # 115809) to produce the HIV-1 LTR-eGFP virus, or with the pCTS.Luc transfer vector (a gift from Stéphane Emiliani) to produce HIV-1 LTR-CMV-Luc-LTR virus (with VSV-G).

HIV-1 (Sup. Fig. 2B) and HIV-2 (Fig. 2C) replicative viruses pseudo-typed VSV-G were produced by transfection of pNL4.3 WT or pGLAN WT (gift from Akio Adachi(69)) and VSV-G vectors and titered on HeLa TZMbl reporter cells. HIV-2 ΔEnv-luciferase viruses, deleted of the Env gene and with luciferase in place of Nef, were obtained following transfection of pGLANΔEnv luciferase (gift from Akio Adachi(70)) and VSV-G vectors.

### Cell lines

#### Cell lines

HeLa (CCL-2) and HEK293T (CRL-3216) cells were cultured in DMEM and Jurkat (TIB-152) and J-Lat A1 cells were cultured RPMI containing 10% heat-inactivated fetal bovine serum (FBS), 1000 units/mL penicillin, and 1000 µg/mL streptomycin.

HeLa KO CTRL, KO TASOR and KO MORC2 cells were generated by the transduction of HeLa cells by VLP containing CRISPR-sgRNA-Cas9 construct targeting TASOR or MORC2 genes. Three days after transduction, these cells were treated by puromycin (1mg/mL) during 5 days. After selection, these cells were diluted and plated in 96-well plates in order to obtain a single cell per well to make a clonal selection. After clonal amplification, MORC2 and TASOR protein levels in each different clone were checked by western blot.

The HIV-1 latency model J-Lat A1 was generated and described by Jordan et al. J-Lat A1 KO CTRL and MORC2 KD cells were generated by transfection of pLentiCRISPRv2-sgCtrl-Cas9 and pLentiCRISPRv2-sgMORC2-Cas9 respectively, with DMRIEC reagent. Transfected cells were cultured for 3 days prior to puromycin selection (1µg/mL during 3 days). The J-Lat A1 MORC2 KD cells were sorted by flow cytometry with the BD FACS ARIA3 cytometer to separate three populations of cells: GFP-negative cells, GFP-positive-cells with low GFP expression (Dim) and GFP-positive cells exhibiting high GFP expression (Bright). Cells were then amplified for two weeks in culture before subsequent experiments.

HeLa LTR-ΔTAR-Luc cells were generated in the laboratory of Stéphane Emiliani from the HeLa LTR-Luc cells described in(71)

The polyclonal Latent HIV-1-eGFP Jurkat model was generated and described in (Matkovic 2022). These cells were transfected by pLentiCRISPRv2-sgCtrl-Cas9 and pLentiCRISPRv2-sgMORC2-Cas9 with DMRIEC reagent in order to generate HIV-1-LTR-eGFP KD Ctrl or KD MORC2 Jurkat cells respectively. PBMCs from the blood of anonymous donors (obtained in accordance with the ethical guidelines of the Institut Cochin, Paris and Etablissement Français du Sang) were isolated by Ficoll (GE Healthcare) density-gradient separation. CD4+ T cells were isolated by positive selection using magnetic CD4 human MicroBeads (Miltenyi Biotec). Cells were activated with CD3/CD28 agonists (T Cell transact) and stimulated with human IL-2 (50U/mL) for 3 days. Cells were then washed twice with Dulbecco’s PBS 1x and lysed with RIPA buffer to assess TASOR and MORC2 protein levels by Western-Blot.

### siRNA treatment

HeLa cells were transfected by siRNA with DharmaFECT1 (Dharmacon, GE Lifesciences). The final concentration for all siRNA was 100 nM. All siRNAs used are listed in Table S1.

### Luciferase activity assays

Cells were washed twice with PBS 1x then lysed with the lysis buffer cell culture Lysis Reagent 1X (Promega) and centrifugated 10 min at 15000g. Luciferase activity was measured using a luciferase assay system from Promega and a TECAN multimode reader Infinite F200 Pro. Luciferase activity data was normalized on protein concentration with the use of Pierce BCA Protein Assay Kit (232255, Thermo-Fisher).

### Western-Blot and antibodies

For all lysates, proteins were separated through 4-12 % Bis-Tris gradient gels, then transferred onto PVDF membrane (ThermoFisher: 88520) and revealed by immunoblot. Antibodies (in 5% skimmed milk in PBS-Tween 0.1%) are listed in Table S2. All secondary antibodies anti-mouse (31430, lot VF297958, ThermoFisher) and anti-rabbit (31460, lots VC297287, UK293475 ThermoFisher) were used at a 1/10000 dilution before reaction with Immobilon Forte Western HRP substrate (WBLUF0100, Merck Millipore). Signals were acquired with Fusion FX (Vilber).

### Flow cytometric analysis

J-Lat A1 and Jurkat cells were washed with PBS and resuspended in PBS-EDTA 0,5 mM. Data was acquired with a BD Accuri C6 cytometer and the FlowJo V10 software. At least 10,000 events in P1 were collected, the GFP-positive population was determined using a GFP-negative population when possible (for Jurkat cells) or arbitrary (as for J-Lat cells) and the same gate was maintained for all conditions. Analysis was performed on the whole GFP-positive population.

Regarding reactivation assay, Jurkat cells are reactivated with 1 ng/mL of TNF-α.

### Chromatin immunoprecipitation

HeLa LTR-ΔTAR-Luc cells were grown in 10 cm dishes. 10×106 cells for each condition were dual cross-linked with 2mM EGS (ethylene glycol bis (succinimidyl succinate)) (#21565 ThermoFisher) at room temperature (RT) for 40min, washed twice in PBS and then fixed in 1% formaldehyde (F8775 Sigma) at room temperature (RT) for 10min and then quenched with glycine 0.125M (7005S Cell Signaling) at RT for 10min. Cells were then washed twice in PBS and scraped into a new tube. After centrifugation at 300xg, 4min, 4°C, pellets were lysed twice in 1mL NP-40 lysis buffer (10mM Tris-HCl pH 7.5, 10mM NaCl, 3mM MgCl2, 0.5% NP-40) centrifuged at 500xg, 4min, 4°C, in order to fraction the cells and keep only the nuclei. Nuclei were then resuspended in 250µL of Sonication buffer (50mM Tris-HCl pH 8.0, 10mM EDTA, 1% SDS) with an anti-protease cocktail (A32965, ThermoFisher) and sonicated for 8 minutes at 4°C (10s ON, 50s OFF) with a Diagenode BIORUPTOR Pico. The sonicates were then pooled and left to chill on ice for 30min in order to let the SDS precipitate. After centrifugation at 16000xg, 10min, 4°C, the supernatant is collected in a new tube. The purified sonicate is then equilibrated with 1.4 vol of TE buffer, 0.026 vol of Triton X-100 (T8787 Sigma), 0.026 vol of 10% sodium deoxycholate and anti-protease cocktail. Then 100µL are collected in a new tube as the input chromatin fraction. Meanwhile, 26µL of Protein A beads (10002D, Invitrogen) were washed twice with 1mL of Bead wash buffer (20mM Tris-HCl pH 8.0, 150mM NaCl, 2mM EDTA, 1% Triton X-100) and once with BSA 5% in PBS on magnetic rack and then resuspended in 500µL of BSA 5% and 5µg of anti-IgG or anti-MORC2 antibodies (#A300-149A, Fortis Life Science) before being wheeled for 2h at RT. Then beads were isolated on a magnetic rack, to remove the supernatant and the remaining purified sonicate was split between each condition before being wheeled overnight at 4°C. The next day, beads were washed twice with a Low Salt buffer (SDS 0.1%, Triton 1%, 2mM EDTA, 20mM Tris HCl pH 8, 150mM NaCl), twice with a High Salt buffer (SDS 0.1%, Triton 1%, 2mM EDTA, 20mM Tris HCl pH 8, 500mM NaCl), once with a LiCl buffer (0.25M LiCl, NP-40 1%, 1mM EDTA, 10mM Tris HCl, 1% Deoxycholate) and once with TE buffer. Beads were then eluted with 170µL of Elution buffer (10mM Tris-HCl pH 8.0, 1mM EDTA, 1% SDS) for 30min at 65°C; the eluate was transferred to a new tube. Then, 5µL of Proteinase K (EO0491, ThermoFisher) was added for each condition and inputs and incubated overnight at 55°C. DNA was then purified by adding 200µL of Phenol/Chloroform/Isoamyl alcohol (25:24:1) (327115000, ThermoFisher), vortexing and centrifuging at maximum speed for 15min at RT. The upper phase was then transferred into a new tube and 0.5 volume of NH4OAc, 1µL of glycogen and 2.5 volume of 100% ethanol were added to precipitate DNA; then, after vortexing, tubes were incubated at -80°C for 1h. After 30min of centrifugation at maximum speed at 4°C, two wash steps were performed with 1mL of ethanol 70%. Pellets were air-dried for 5min before resuspension in 80µL of DNAse free water. qPCR was finally performed as described in the RT-qPCR section.

### Nuclear Run On (NRO)

HeLa LTR-ΔTAR-Luc cells were plated in 10 cm dishes at 2,5 billion cells and the following day transfected with siRNA CTRL, TASOR + CTRL and MORC2 + CTRL at a final concentration of 100 nM each. NRO was performed as precisely described by Roberts and colleagues(72), except for these specific points: At step 14, RNA extractions with rDNase treatment were performed with NucleoSpin RNA, Mini kit (740955.250, Macherey-Nagel). At Step 44, Reverse transcription was performed with Maxima First Strand cDNA Synthesis Kit with dsDNase (K1672, ThermoFisher). qPCR was performed as described in the RT-qPCR section.

### RT-qPCR

RNA extractions were performed through NucleoSpin RNA, Mini kit (740955.250, Macherey-Nagel). Reverse transcription steps were performed with Maxima First Strand cDNA Synthesis Kit with dsDNase with 1µg of total RNA. qPCRs were performed with a LightCycler480 (Roche) using a mix of 1x LightCycler 480 SYBR Green I Master (Roche) and 0.5 μM primers. The sequences of 5’-3’ oriented primers are listed in Table S2.

### Cell fractionation, Immunoprecipitation and Western-Blot

All immunoprecipitation experiments are performed in the nuclear fraction of HeLa cells (WT, KO CTRL, KO MORC2 and KO TASOR). Cells grown in 10 cm dishes were washed with PBS 1X. After trypsinization, cells were recovered in 1.5 mL tubes and washed once with ice-cold PBS. After 4 min of centrifugation at 300g, 1 mL of Cytoplasmic Lysis Buffer

(10 mM TRIS-HCl pH7.5, 10 mM NaCl, 3 mM MgCl2, and 0.5% IGEPAL® CA-630 (I8896-100ML-Merck)) was added on the cell pellet and resuspended pellet was incubated on ice for 5 min. Cells were then centrifuged at 300g for 4 min at 4 °C. The supernatant was saved for cytoplasmic fraction. The pellet was washed with 1 mL of Cytoplasmic Lysis Buffer and centrifuged at 300g for 4 min at 4 °C twice. Finally, the nuclear pellet was lysed with 300 μL of RIPA Buffer. 250 μg of nuclear proteins was wheeled with 3μg of anti-IgG antibody or 3μg of anti-CNOT1 antibody or 3μg of anti-MORC2 or 3μg of anti-DDK at 4°C. The following day, pre-washed PierceTM Protein A/G Magnetic Beads (88802, ThermoFisher) were added to the samples for 1h at room temperature. The captured beads were washed 3 times with the wash buffer (150mM NaCl, 50mM Tris pH7,5). Finally, elution was performed by adding 1x Laemmli solution containing DTT 0,1M and heated for 10 min at 95°C. Western-Blot procedure is described in SI appendix.

## Data and Reagent availability

Data for sequence analyses (codon alignments, and phylogenetic tree) are available at https://doi.org/10.6084/m9.figshare.22337539, https://doi.org/10.6084/m9.figshare.22337542, and https://doi.org/10.6084/m9.figshare.22337545. All other data and reagents are available from the corresponding authors.

## Acknowledgements

We thank all the members of the Retrovirus, Infection and Latency team and Stéphane Emiliani for discussions. We also thank the members of the LP2L team at CIRI, Lyon, for support. We acknowledge the CYBIO and GENOM’IC platforms of the Institut Cochin.

This work was supported by grants from SIDACTION, the French Research Agency on HIV and Emerging Infectious Diseases ANRS/MIE and Fondation pour la Recherche Médicale (FRM, PROJECT EQU202203014684 attributed to F.MG.). R.M. was supported by Sidaction and FRM; M.M.M. by SIDACTION ; An.L., S.M. and P.L. by Université de Paris Cité ; V.V. by ANRS. This work in the laboratory of L.E. is supported by grants from the ANRS/MIE (#ECTZ118944 to LE) and Sidaction (n°21-1-AEQ-12972-2 to LE and FMG). Al.L is supported by a PhD fellowship from Sidaction (2020 - n°12673).

---

**Supplementary Figure 1:**
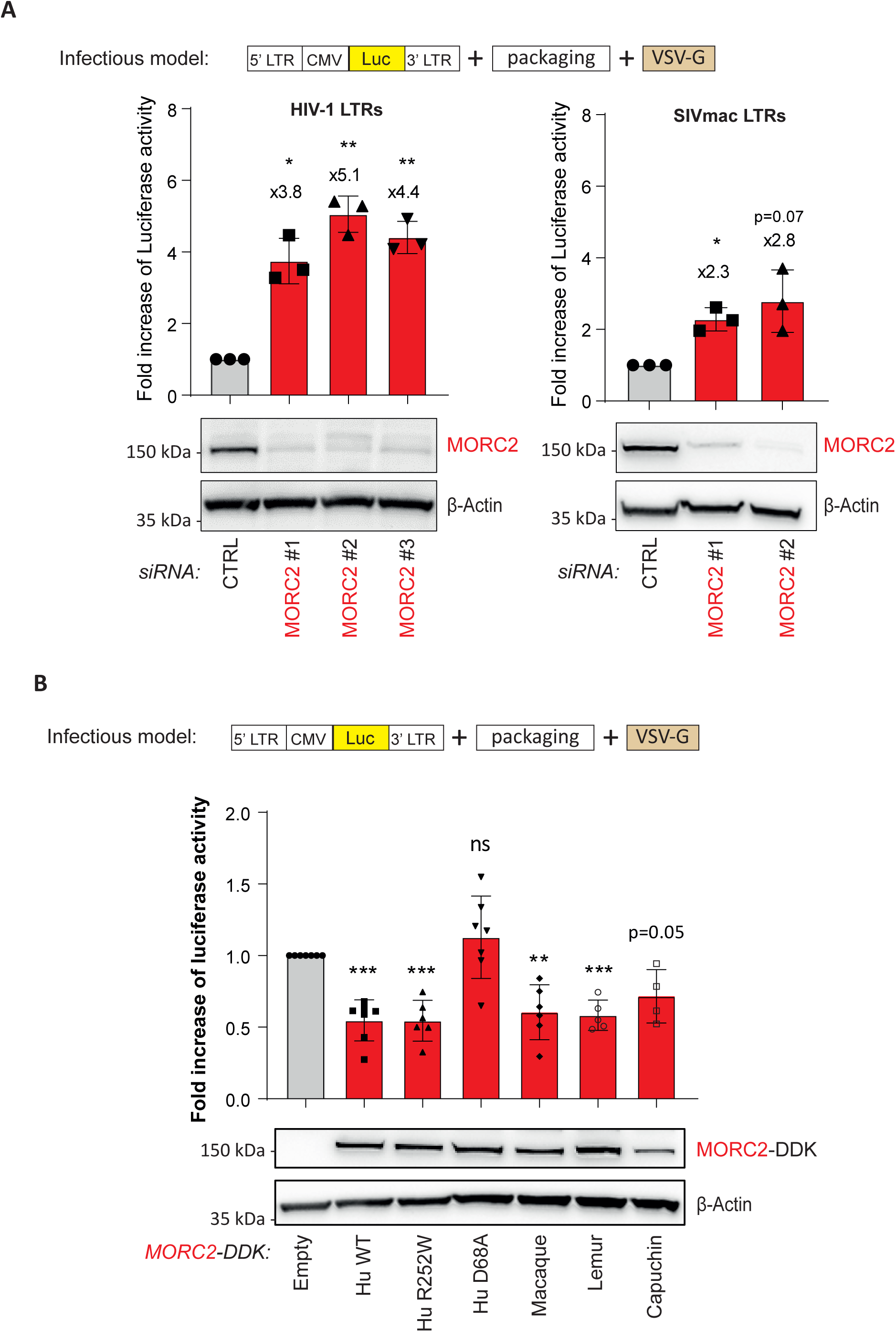
**A. Depletion of MORC2 favors HIV-1- and SIVmac-derived virus expression.** HeLa cells were infected by HIV-1- or SIVmac-derived viruses expressing luciferase (luc) (LTR-CMV-Luc viruses), then the following day, treated during 72h by siRNA CTRL or three (or two) different siRNA MORC2. Luc gene expression was measured by Luc activity assay (*n* = 3; each replicate is presented along with the mean values and the SD. Lognormality has been verified with a Shapiro-Wilk test prior to a One sample t test with a theoretical mean of 1. \**p* < 0.05 \*\**p* < 0.01) **B. Overexpression of Human and orthologous non-Human primate MORC2 proteins induces a decrease of HIV-1-derived virus expression in Human cells.** HeLa KO MORC2 cells were infected by HIV-1 LTR-CMV-Luc viruses, then the following day, transfected by vectors expressing different MORC2 proteins (four orthologous proteins from divergent primate species: human (Hu), rhesus macaque (Mac), Northern greater galago (Lem), white-faced capuchin (Cap)); MORC2 D68A, an ATPase defective protein, used as a control; and MORC2 R252W, which is the protein produced by a mutated *MORC2* gene in the Charcot Marie-Tooth disease and which is supposed to hyperactivate HUSH. Luc gene expression is measured by Luc activity assay (Minimum *n* = 4; each replicate is presented along with the mean values and the SD. Lognormality has been verified with a Shapiro-Wilk test prior to a One sample t test with a theoretical mean of 1. \**p* < 0.05 \*\**p* < 0.01 \*\*\**p* < 0.001)

**Supplementary Figure 2:**
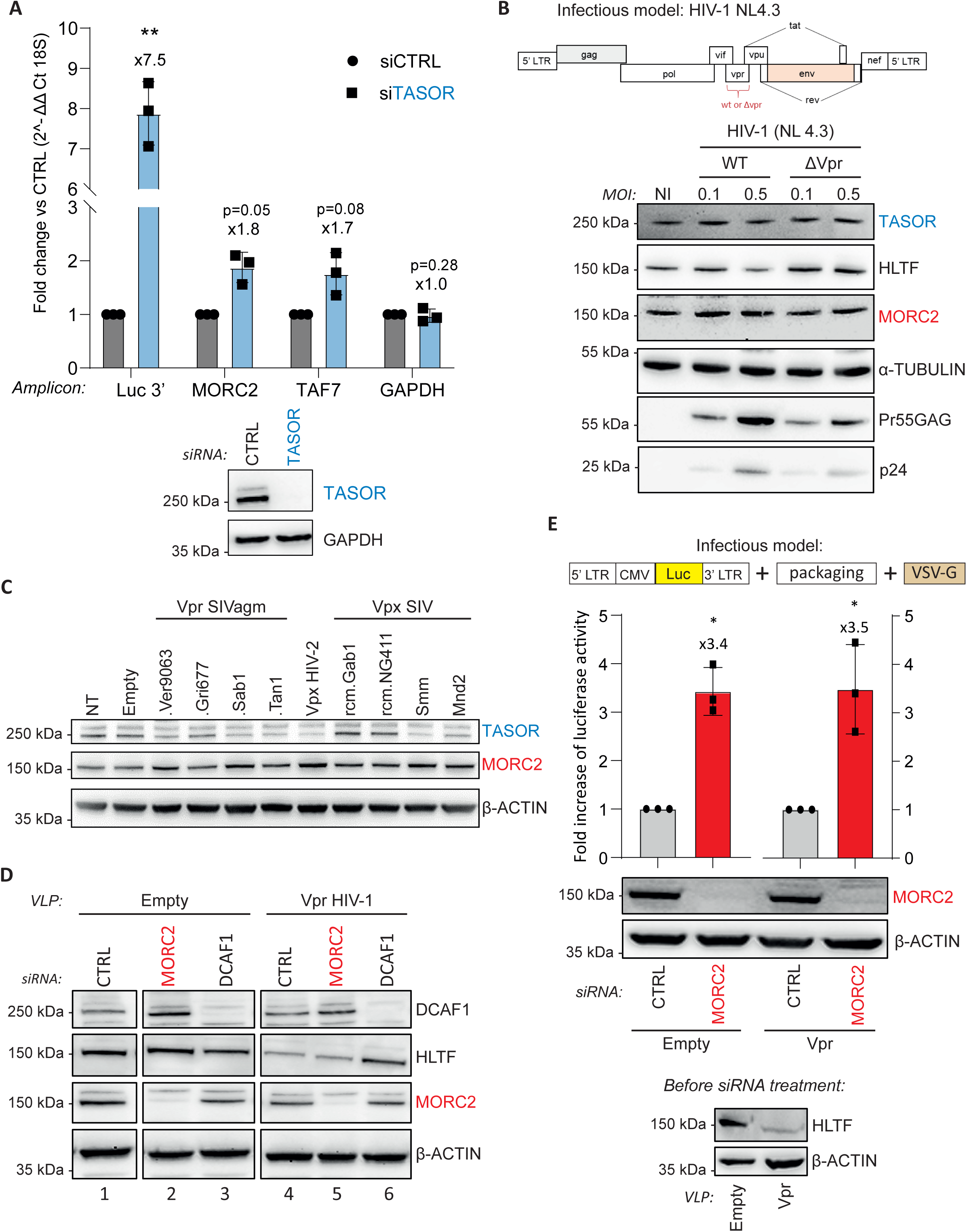
**A. TASOR depletion induces an increase of MORC2 mRNA level.** HeLa HIV-1 LTR ΔTAR-Luc cells were transfected by Ctrl or TASOR siRNA and, 72h post-transfection, cells were lysed and RT-qPCR was performed to analyze mRNA MORC2 levels (*n* = 3; each replicate is presented along with the mean values and the SD. Lognormality has been verified with a Shapiro-Wilk test prior to a One sample t test with a theoretical mean of 1. \*\**p* < 0.01). **B. HIV-1 does not induce MORC2 protein depletion.** Jurkat T cells were non-infected (NI) or infected with complete HIV-1 viruses (WT) or HIV-1 viruses deleted of Vpr (ΔR), at two different MOI (0.1 and 0.5). Indicated proteins are revealed by western blot. HLTF protein is a positive control of depletion by Vpr-containing HIV-1. **C. Vpr and Vpx proteins from divergent primate lentiviruses do not impact MORC2 expression**. Viral proteins from divergent origins were delivered by VLP: Vpr from SIVagm.Vervet (ver), Grivet (gri), Sabaeus (sab), Tantalus (tan), Vpx from SIVsmm (infecting sooty mangabey), SIVmnd-2 (mandrill), SIVrcm.gab1 and SIVrcm.ng411 (2 strains infecting red-capped mangabey). **D. MORC2 is not required for HIV-1 Vpr-mediated HLTF degradation.** HeLa cells were treated by MORC2 siRNA, then transduced by VLP containing HIV-1 Vpr. Cells were lysed the day after, and western-blot performed. **E. MORC2 remains active following HIV-1 Vpr treatment.** HeLa cells were infected by the HIV-1 LTR-CMV-Luc virus, 2 days later, HIV-1 Vpr was delivered thanks to VLP transduction, next day, MORC2 was depleted by siRNA. Luciferase gene expression was measured by Luciferase activity assay (*n* = 3; each replicate is presented along with the mean values and the SD. Lognormality has been verified with a Shapiro-Wilk test prior to a One sample t test with a theoretical mean of 1. \**p* < 0.05)

**Supplementary Figure 3:**
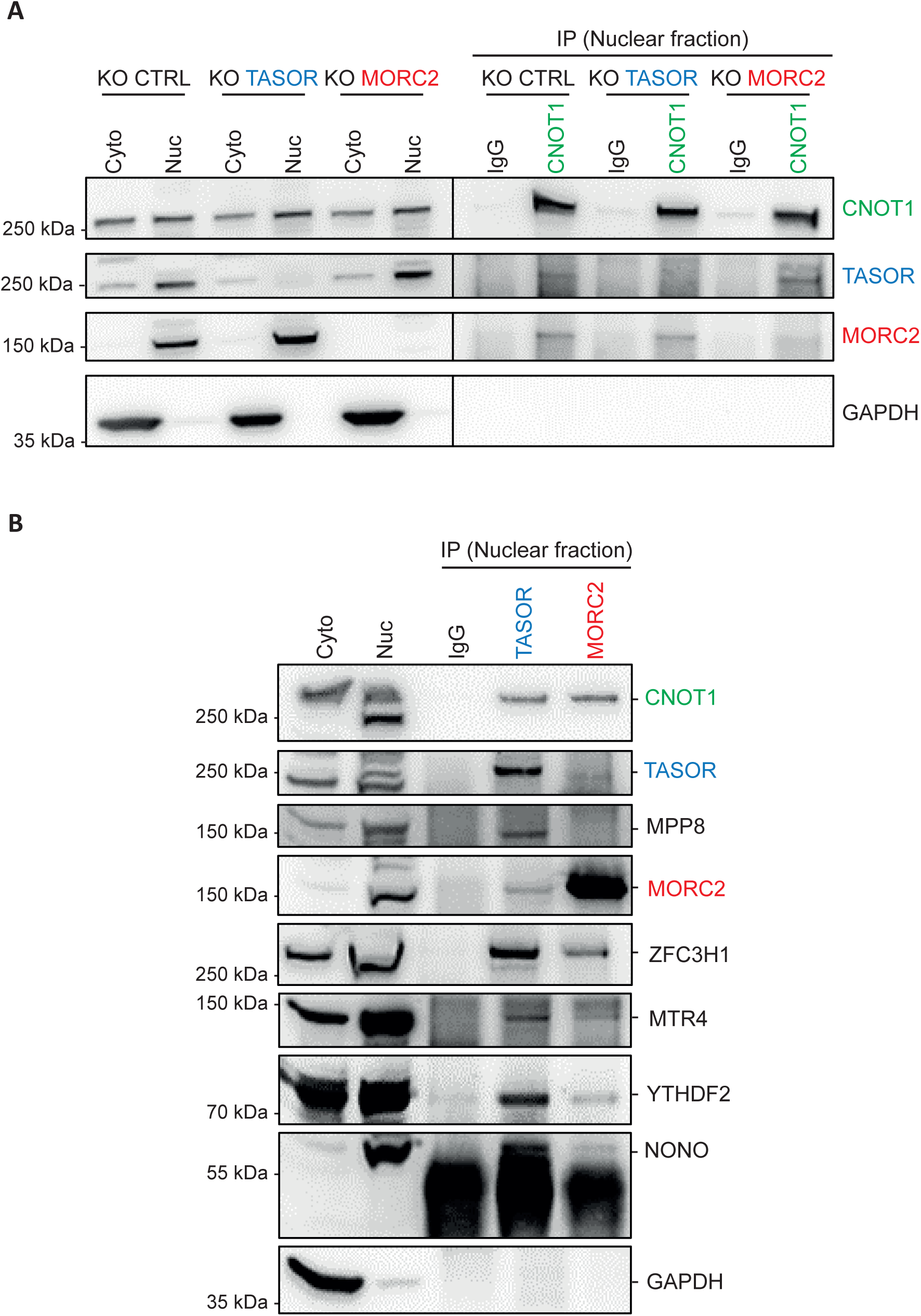
**A. CNOT1 interacts with MORC2 in the absence of TASOR, and with TASOR in the absence of MORC2, in the nucleus of HeLa cells.** Nuclear fraction of Hela KO CTRL, KO MORC2 and KO TASOR cells was isolated. Endogenous CNOT1 was immunoprecipitated with an antibody specific to CNOT1 and Western-Blot was performed. GAPDH is a negative control for nuclear fraction isolation. **B. Interaction of endogenous TASOR or MORC2 with RNA metabolism proteins**. Nuclear fractions of HeLa cells were isolated and TASOR or MORC2 were immunoprecipitated with specific antibodies, then western blot was performed. GAPDH is a negative control and NONO a positive control for nuclear fraction isolation.

**Supplementary Figure 4:**
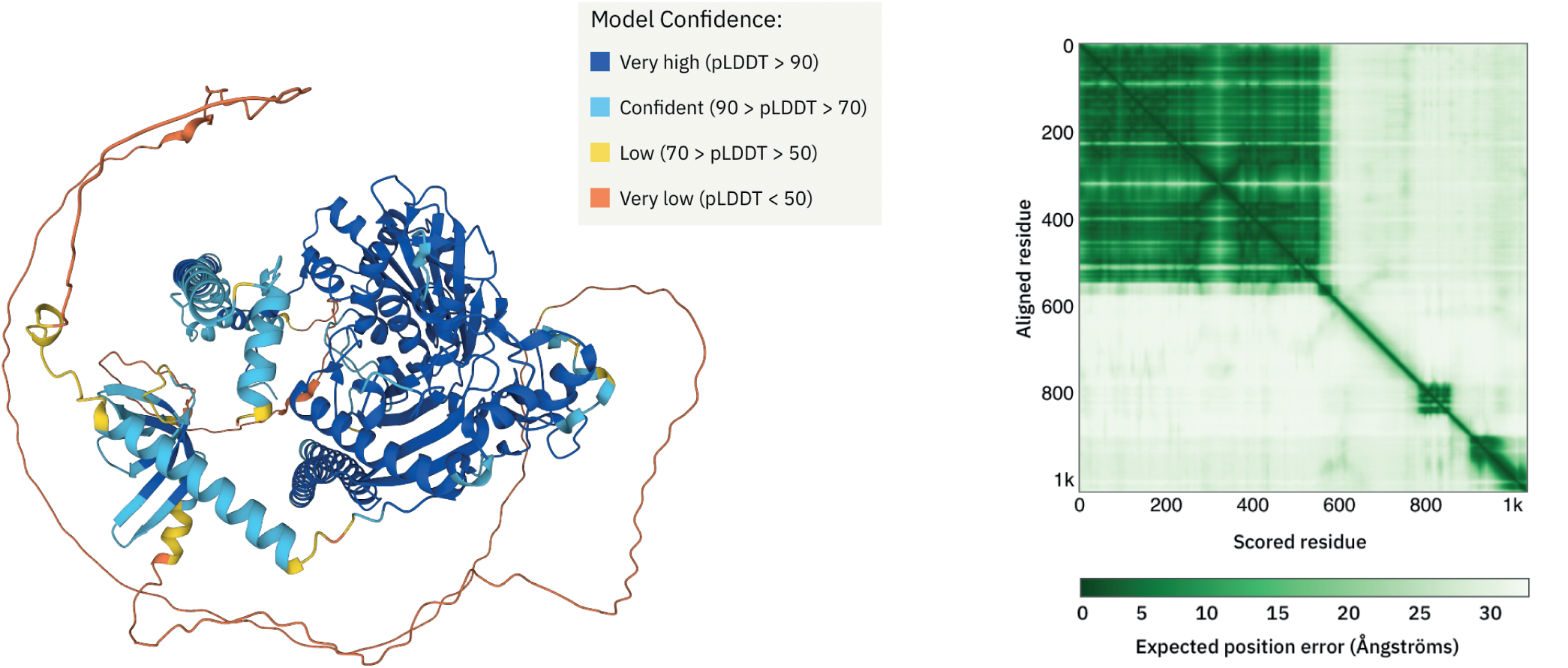
Model confidence of MORC2 AlphaFold predicted structure. *Left*. Model confidence of MORC2 AlphaFold predicted structure. pLDDT (predicted local distance difference test): Per-residue confidence score. *Right*. Predicted aligned error plot of MORC2 AlphaFold predicted structure. Dark green represents low distance error in Ångströms, light green represents high distance error. Figures are from AlphaFold Protein Structure Database website.

**Supplementary Figure 5:**
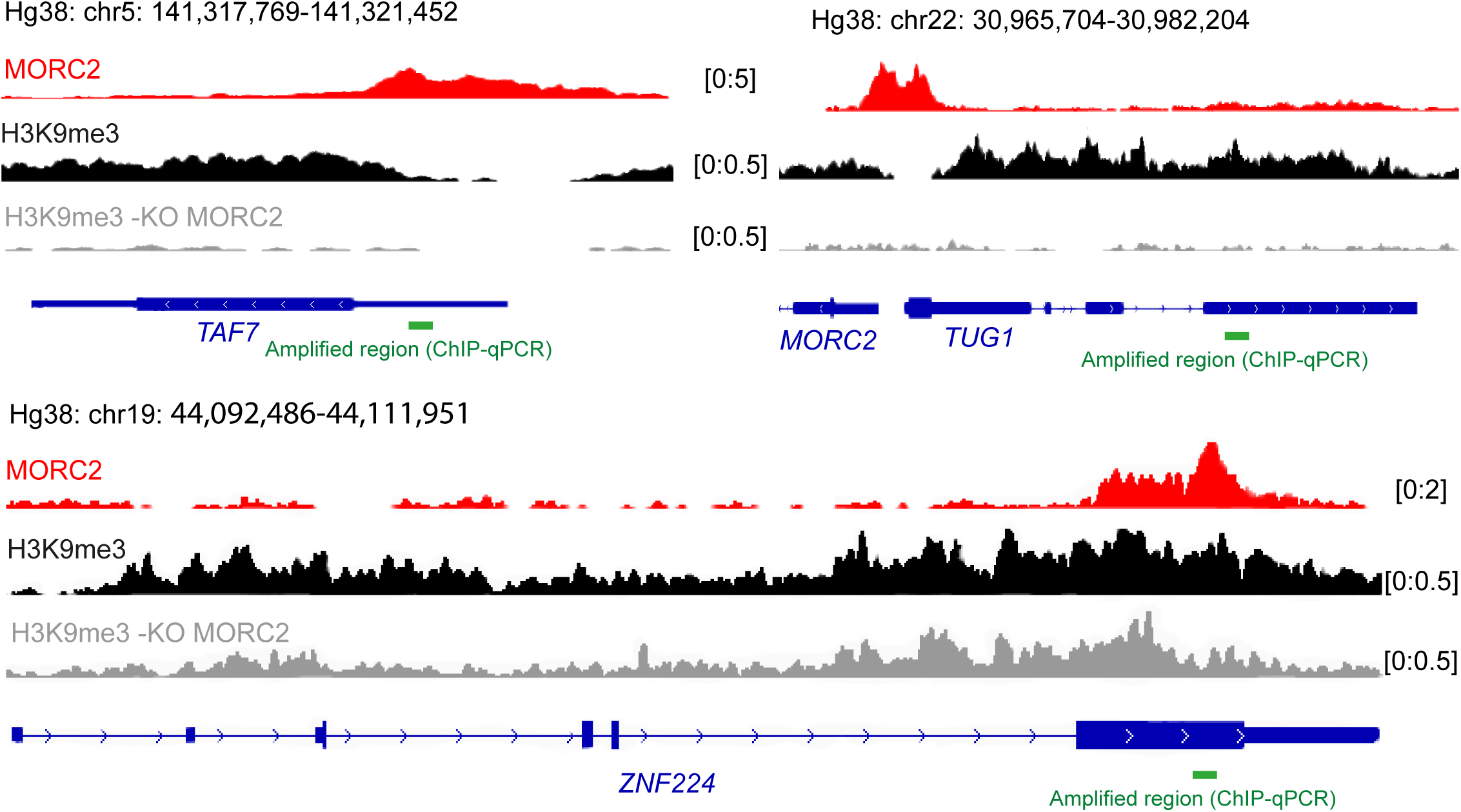
MORC2 binds the genomic locus of TAF7, TUG1 and the last exon of ZNF224. IGV profiles of MORC2, H3K9me3 (WT and MORC2 KO) performed in K562 cells by (Liu 2018) and available at GEO under accession number GSE95374. The amplified region of our ChIP qPCR analyses is denoted in green. These targets stand as positive controls of MORC2 ChIP-qPCR in our study. The sequences of the primers are listed in Table S2.

**Supplementary Table 1.**
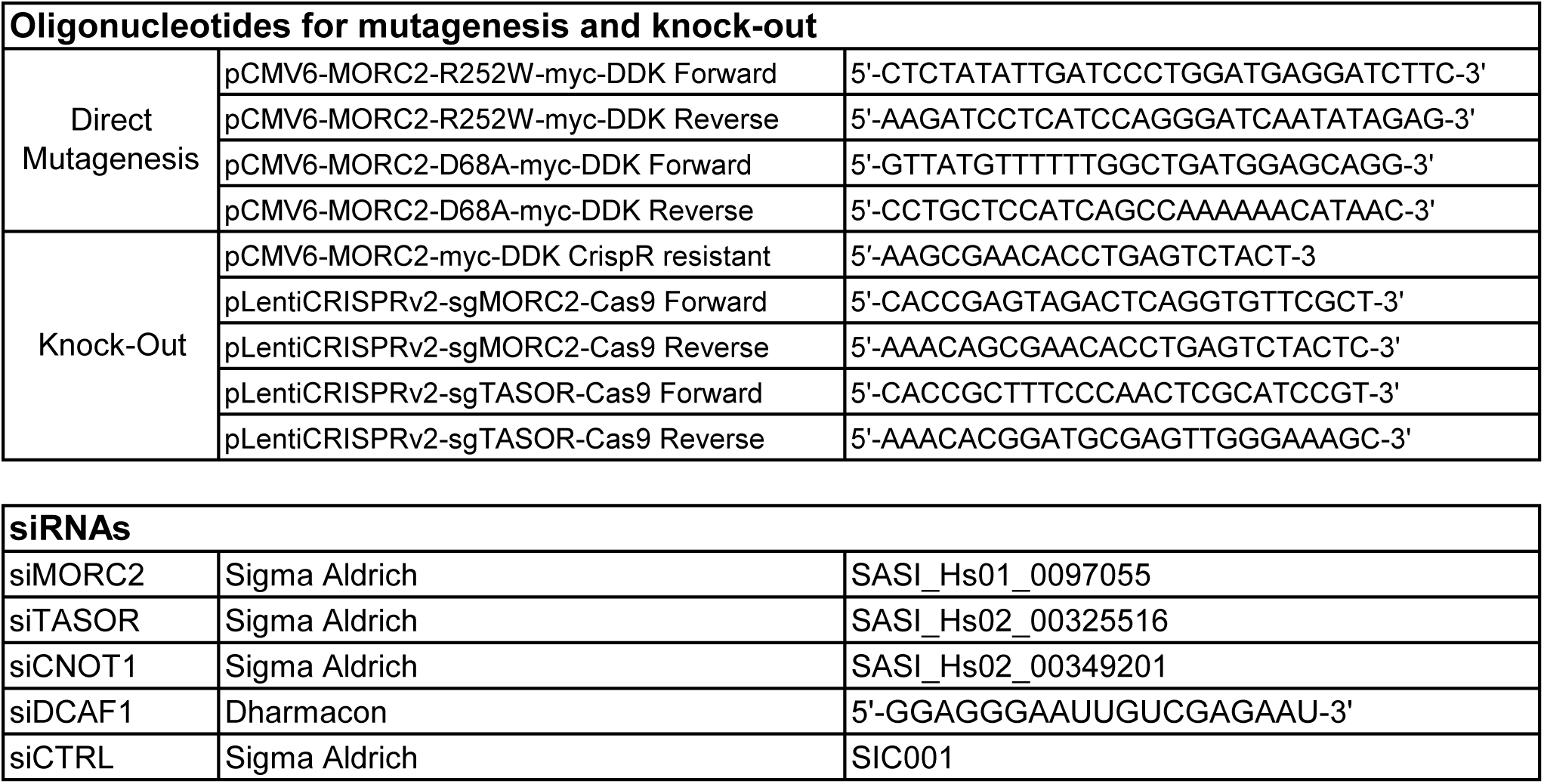

**Supplementary Table 2.**
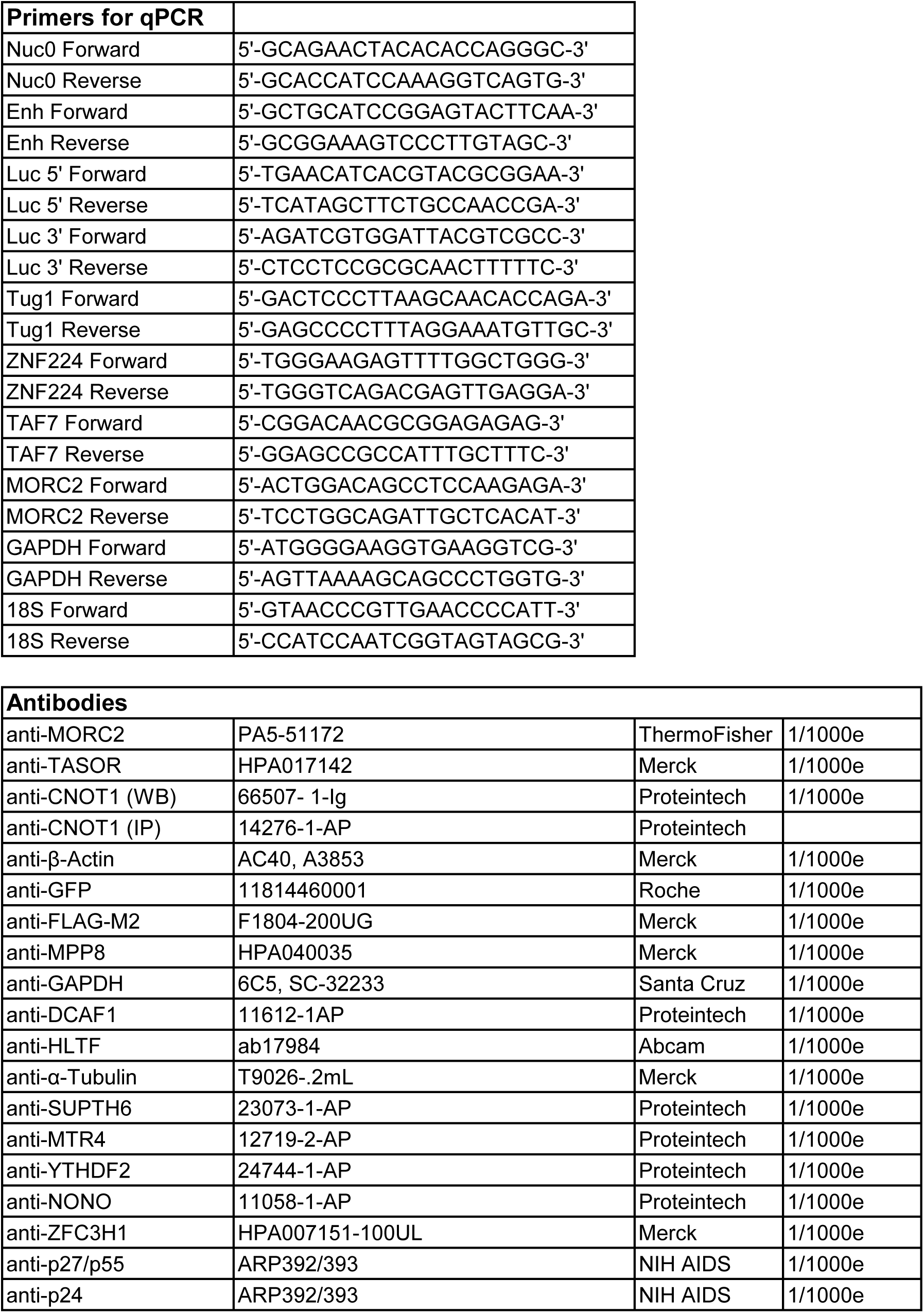

